# Predicting HIV Incidence in the SEARCH Trial: A Mathematical Modelling Study

**DOI:** 10.1101/376244

**Authors:** Britta L. Jewell, Anna Bershteyn

## Abstract

**Introduction:** SEARCH is one of four randomized-controlled trials (RCTs) investigating the strategy of community-based treatment-as-prevention (TasP) for the reduction of HIV incidence in sub-Saharan Africa. SEARCH takes place among 32 pair-matched rural communities in three regions of East Africa and exceeded the UNAIDS 90-90-90 targets for HIV testing, linkage to care, and viral suppression in the intervention arm. We used mathematical modeling to estimate expected 3-year cumulative HIV incidence in both arms of the trial, using different assumptions about two main sources of uncertainty: scale-up of antiretroviral therapy (ART) in the control arm, and the degree of mixing between SEARCH residents and non-residents.

**Methods:** We used the HIV modelling software EMOD-HIV to configure and calibrate a new model of the SEARCH communities. The 32 trial communities were clustered into six nodes (three for the control arm and three for the intervention arm) using k-means clustering based on community HIV prevalence, male circumcision rates, mobility, and geographic region. The model was parameterized using data on demographics, HIV prevalence, male circumcision rates, and viral suppression data collected at trial baseline in 2013, and calibrated to node-specific and age-specific HIV prevalence, ART coverage, and population size. Using data on ART scale-up in subsequent follow-up years in the trial, we varied linkage to ART in the control arm and the degree of external mixing between SEARCH residents and non-residents.

**Results:** If no external mixing and no additional control arm ART linkage occurred, we estimate the trial would report a relative risk (RR) of 0.60 (95% CI 0.54-0.67, p<0.001), with all simulations showing a significant result. However, if SEARCH residents mixed equivalently with non-residents and ART linkage in the control arm also increased such that the control arm also exceeded the 73% viral suppression target, the RR is estimated to be 0.96 (95% CI 0.87-1.06, p=0.458) and 72% of simulations are non-significant. Given our “best guess” assumptions about external mixing and year 3 data on ART linkage in the control arm, the RR is estimated to be 0.90 (95% CI 0.81-1.00, p=0.05), with 49% non-significant simulations.

**Conclusion:** The SEARCH trial is predicted to show a 4-40% reduction in cumulative 3-year incidence, but between 18-72% of simulations were non-significant if either or both ART linkage in the control arm and external mixing are substantial. Despite achieving the 90-90-90 targets, our “best guess” is that the SEARCH trial has an equal probability of reporting a non-significant reduction in HIV incidence as it does a significant reduction.

## Introduction

In 2011, the HPTN 052 trial reported that antiretroviral therapy (ART) was associated with a 96% reduction in HIV transmission [1], confirming the results of numerous observational studies [2, 3]. While treatment-as-prevention (TasP) has been shown to drastically reduce HIV transmission on an individual level, it has not yet been shown to reduce population-level incidence in the context of a randomized-controlled trial (RCT). However, mathematical modeling and observational studies have strongly suggested that increased ART coverage could have a substantial effect on reducing new HIV infections, among other benefits [4, 5]. In light of this, UNAIDS released the “90-90-90 targets” in 2014, which aim to end the epidemic by achieving 90% testing coverage among HIV-positive individuals, 90% ART linkage for infected individuals, and viral suppression in 90% of those on ART by the year 2020 [6].

The SEARCH (Sustainable East Africa Research in Community Health) Study (NCT01864683) is one of four major randomized controlled trials (RCTs) investigating the strategy of community-based treatment-as-prevention (TasP) for reduction of HIV incidence in sub-Saharan Africa. The SEARCH trial is based in 32 rural communities of approximately 10,000 people each in three regions of East Africa – Eastern Uganda, Western Uganda, and Western Kenya. Eastern Uganda communities are characterized by average HIV prevalence among adults of 4%, while in Western Ugandan and Kenyan communities, adult prevalence averages 7% and 20%, respectively [7]. Phase I of the study, which took place from 2013 to 2016, is investigating the effect of universal HIV treatment and streamlined care on HIV incidence, mortality, and non-communicable disease control, among other outcomes.

At baseline, a census was conducted among all communities to enumerate all individuals in study communities. During the census, individuals were classified as “resident” (i.e., those spending at least 6 months of the past year in the community) or “non-resident” (i.e., those spending at least 6 months of the past year outside of the community). The incidence cohort for Phase I of the study was defined as all resident adults ≥15 years old at baseline. At baseline in 2013, two-week community-wide testing campaigns (CHCs) were conducted in all control and intervention arm communities. At each CHC, all participants were offered multi-disease services, including HIV, hypertension, and diabetes treatment, as well as malaria treatment and deworming for children. Residents who did not attend a CHC were tracked to their homes and offered home-based testing (HBT). Through a combination of CHCs and HBT, SEARCH achieved 89% baseline testing coverage across both arms of the trial at baseline [8].

In the control arm, individuals were offered standard-of-care ART start according to national guidelines. Over the three years of study follow-up, guidelines changed from CD4 counts ≤350 cells/μl to CD4 counts ≤500 cells/μl in Kenya and Uganda in July 2014 and December 2013, respectively [9, 10]. Both countries also adopted guideline changes for universal test and treat in July 2016 (Kenya) [11] and November 2016 (Uganda) [12]. In the intervention arm, HIV-positive individuals were offered immediate and universal ART, regardless of CD4 count or other criteria. Annual testing campaigns were held from follow-up years 1 to 3, and streamlined care interventions (e.g. same-day ART start and 3-monthly refills) were also offered to participants. At baseline, 45% of HIV-positive adult residents were virally suppressed and population-level suppression increased to 80% in the intervention arm of the study after two years of the intervention, exceeding the 90-90-90 targets [13].

Three other major trials are testing the population-level effectiveness of TasP on cumulative HIV incidence in sub-Saharan Africa [14]. Thus far, only one study – the ANRS 12249 trial in KwaZulu-Natal, South Africa – has reported complete results, showing no significant difference between the control and intervention arms of the trial [15]. However, linkage from testing to treatment was lower than expected in both arms of the trial; the cascade for the control and intervention arms combined was 92-58-85% for testing coverage, ART linkage, and viral suppression, respectively, falling short of the 90-90-90 targets [15]. Another study, the HPTN 071 (PopART) trial in Zambia, achieved 78% and 87% testing coverage among men and women, respectively after one year [16]. Among known HIV-positives, 74% of men and 73% of women were also linked to ART, but the study has yet to report viral suppression statistics [16]. Finally, the Botswana Combination Prevention Project (BCPP) observed that at baseline, 83% of HIV-positive individuals were diagnosed, 73% were on ART, and 70% were virally suppressed, nearly meeting the UNAIDS targets prior to the intervention [17]. The SEARCH trial is the only one of these four RCTs to reach 90-90-90 in the initial years of the intervention from substantially lower baseline numbers, and may therefore represent the best opportunity to investigate the population-level effect of TasP in the context of an RCT.

Though ART is proven to reduce transmission on an individual level, there are multiple dimensions of uncertainty in a large-scale trial like SEARCH. First, ART scale-up in the control arm of the trial was unknown between baseline and year 3 at the end of the trial, and it is unclear what effect – if any – the baseline testing campaign had on ART linkage and viral suppression. Second, SEARCH communities are not geographically isolated and residents may have had sexual relationships with individuals outside of the study community who were not receiving the interventions, and mobility has also been associated with increased risk of HIV acquisition for both men and women [18–20]. Lastly, heterogeneity in characteristics of individuals who achieved viral suppression could mean that those responsible for the greatest proportion of transmission remain unsuppressed, potentially substantially affecting the impact of reaching the 90-90-90 targets [21].

In this analysis, we use a mathematical model of the SEARCH Study at baseline, and ART and male circumcision data from follow-up years 1-3 of the trial, to estimate anticipated HIV incidence in and relative risk between the control and intervention arms of the trial following completion of Phase I of the trial. Modelers conducting the study were formally blinded to incidence results in the study, allowing the analysis to serve as a test model validity, while also using modelling to infer potential reasons for the trial outcome that would be consistent with other observations made throughout the trial.

## Methods

We used an existing individual-based model of HIV transmission, EMOD-HIV, to configure a new model of the 32 SEARCH communities in Uganda and Kenya. The model is publicly available (https://github.com/InstituteForDiseaseModeling/EMOD) and has been described previously on multiple occasions [21–26]; briefly, it is a stochastic, individual-based network model of heterosexual and vertical HIV transmission that includes age-specific fertility, age- and sex-specific mortality, four types of heterosexual relationships – commercial, transitory, informal, and marital – with age-specific partnership formation, and a time-varying HIV care cascade encompassing different modes of testing, diagnosis, and linkage to treatment.

In the model, the 32 SEARCH communities were categorized into six nodes – three for each arm of the trial – using k-means clustering based on community HIV prevalence at baseline, population age structure, mobility (>1 month of travel in the prior six months), and male circumcision prevalence [27]. The majority of communities were classified according to their region, with two exceptions; Bware, a control community in Kenya with 6% HIV prevalence and 67% prevalence of traditional male circumcision, was classified with lower prevalence communities in Western Uganda, and Kitwe, an intervention community in Western Uganda with 6% HIV prevalence and a younger age structure, was classified with the lowest prevalence Eastern Uganda node.

The model was parameterized using demographic, HIV prevalence, and viral suppression data from the baseline of the SEARCH Study at the midpoint of 2013 [7]. In the intervention arm nodes of the model, ART scale-up was modelled using viral suppression data from follow-up years 1 and 2, with subsequent incorporation of year 3 data. Model calibration figures, showing 250 best-fit trajectories for node-specific HIV prevalence over time and age-specific HIV prevalence at baseline, are shown in Figure 1. Modelers conducting the study were formally blinded to incidence results in the study.

**Figure 1.**
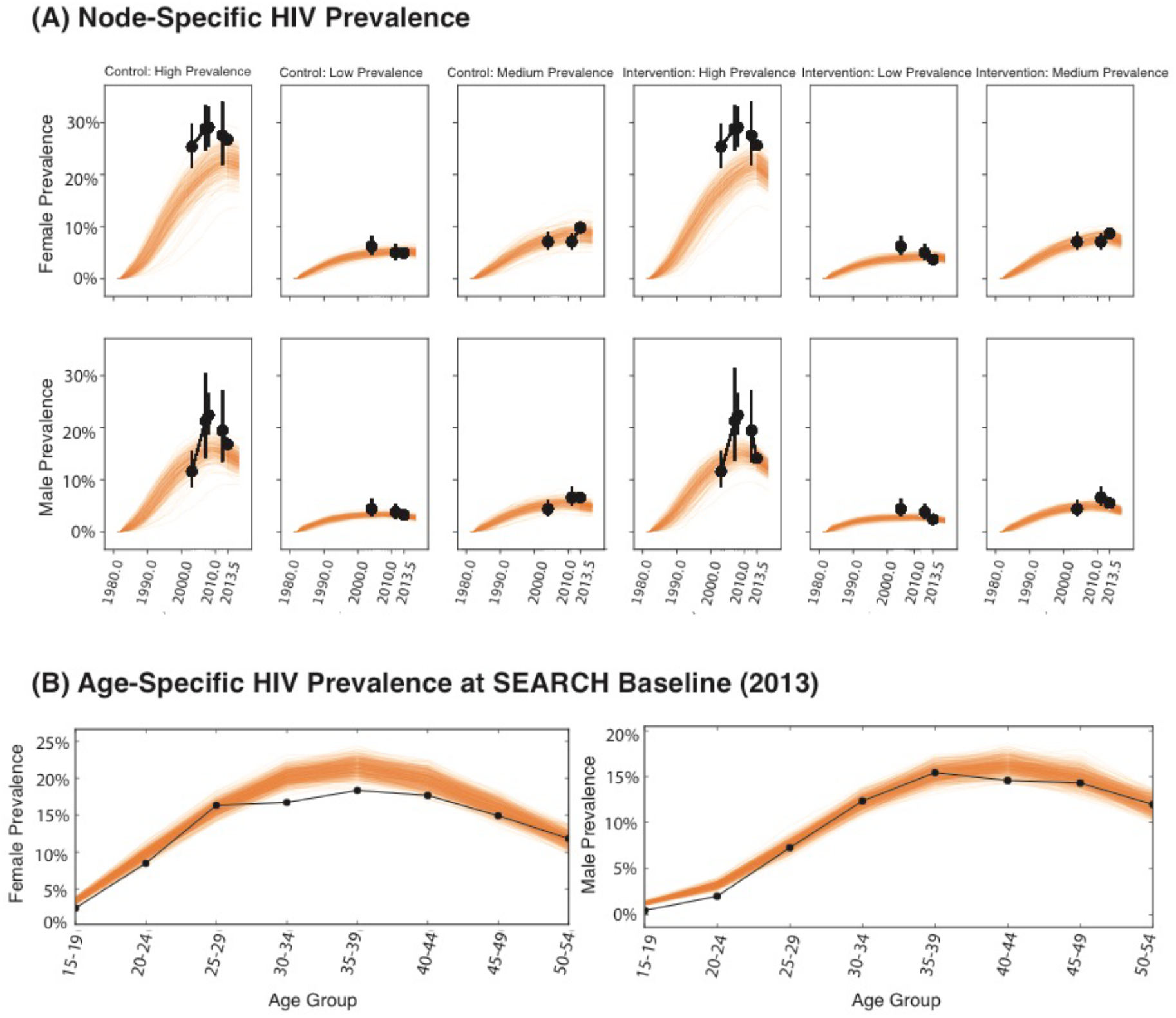
Modeled HIV prevalence among adults (aged 15-49) over time for six nodes (A) and age-specific HIV prevalence at SEARCH baseline in 2013 (B). Each orange line represents one of 250 selected model trajectories; black dots and bars represent data. In (A), data prior to 2013.5 is taken from regional DHS survey data that corresponds with the respective region of nodes of the SEARCH communities, but does not match the boundaries of the communities directly.

We simulated four scenarios across the two greatest sources of uncertainty: ART scale-up in the control arm and external mixing of SEARCH residents with non-residents not receiving the intervention. External mixing is assumed to range from a minimum of zero mixing (i.e., communities are closed cohorts) to a maximum of 1:1 mixing in which SEARCH residents have an equal probability of mixing with a non-SEARCH individual as they do with a SEARCH resident. ART scale-up in the control arm is assumed to depend on the degree to which the baseline testing campaigns resulted in additional linkage to care. We assume the minimum of no additional linkage to result in 69% of HIV-infected individuals in the control communities virally suppressed by the end year 3, and a maximum of 76% of HIV-infected individuals virally suppressed by year 3 resulting from a boost in linkage after baseline CHCs.

Following incorporation of year 3 data, we also predicted “best guess” scenarios based on incorporating viral suppression data in the control arm at the end of year 3, and estimating the prevalence of external mixing using node-specific mobility data from the study baseline. We used the proportion of stable adult residents spending at least one night away from home in the prior month to approximate the availability of mixing with non-SEARCH individuals, and also provide a prediction of the impact of the interventions if no mixing had occurred, given scale-up of ART in the control arm from baseline to year 3. Mobility varied from 25-41% and viral suppression in the control arm was approximately 10% lower than the intervention arm by year 3.

## Results

We projected incidence in the communities in the absence of any baseline testing campaign in the trial or any subsequent SEARCH-specific interventions, but including male-circumcision scale-up and changes in ART eligibility due to national guidelines over time. Due to slightly higher HIV prevalence in the intervention arm of the trial at baseline (10.2% compared to 10.0% in the control arm), three-year cumulative incidence is estimated to be 1.78% in the control arm and 1.83% in the intervention arm in the absence of any interventions. Incidence would be expected to decline from 0.75/100 person-years (PY) to 0.70/100 PY from years 1 to 3 of the trial in the absence of SEARCH interventions, due to scale-up of male circumcision and ART in the normal cascade of care.

Figure 2 shows the density distribution of three-year cumulative incidence for the four extreme points of the sweep across two greatest dimensions of uncertainty: the amount of additional ART linkage in the control arm due to the baseline testing campaign, and the degree of external mixing with non-SEARCH residents. In the first scenario (Figure 2A), we assume no external mixing and no additional linkage (i.e., a closed cohort with no active control); in the second (Figure 2B), we assume no external mixing and maximum linkage due to the baseline testing campaign; in the third (Figure 2C), we assume equivalent mixing between SEARCH residents and non-residents and no additional linkage; in the fourth (Figure 2D), we assume equivalent mixing and maximum additional ART linkage.

**Figure 2:**
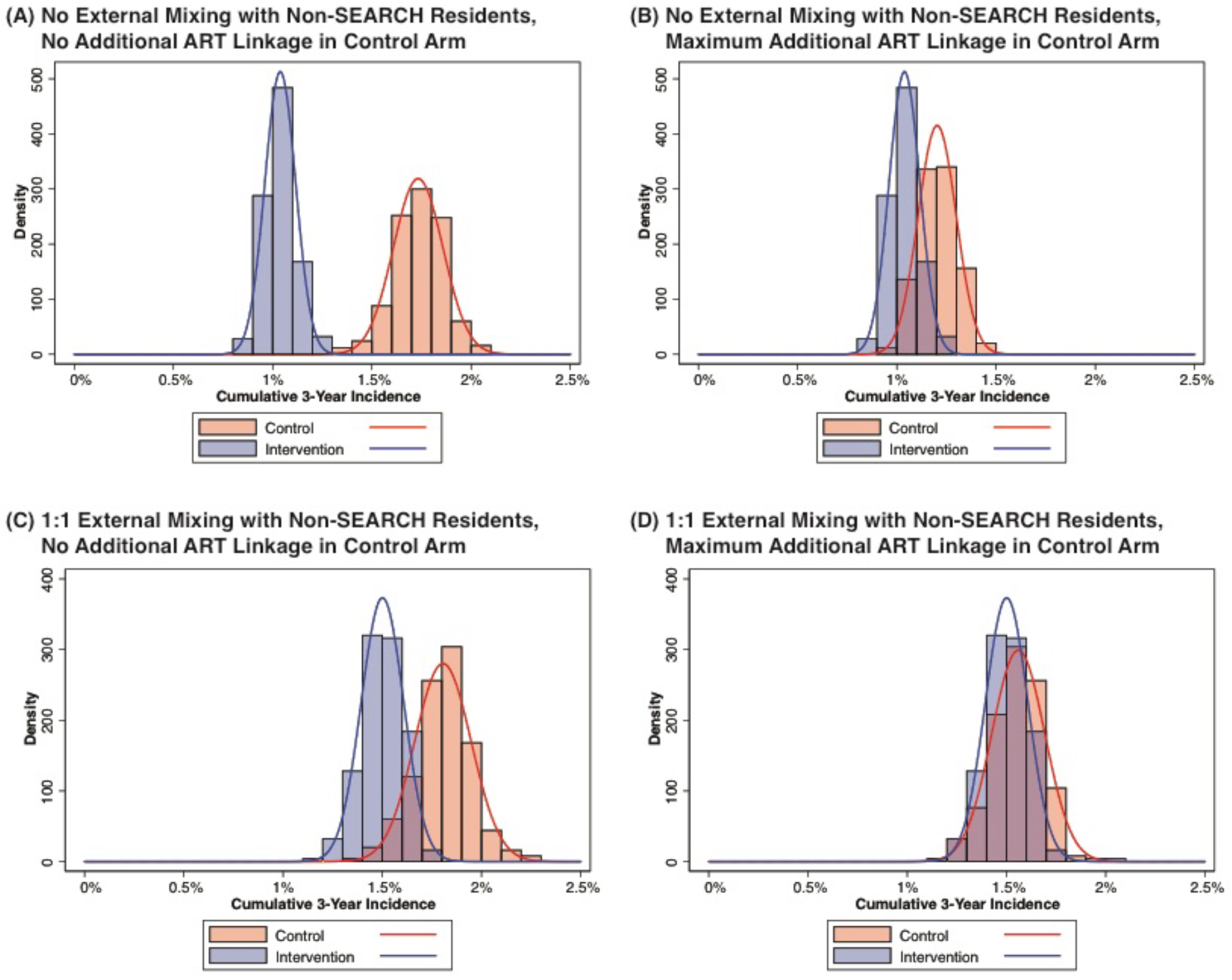
Cumulative 3-year incidence in the control and intervention arms of the SEARCH Study, based on different assumptions of mixing between SEARCH and non-SEARCH residents and ART scale-up in the control arm: (A) no external mixing with non-SEARCH residents and no additional ART linkage in the control arm; (B) no external mixing with nonSEARCH residents and maximum additional ART linkage in the control arm; (C) 1:1 external mixing with non-SEARCH residents and no additional ART linkage in the control arm; and (D) 1:1 external mixing with non-SEARCH residents and maximum additional ART linkage in the control arm.

In the “closed cohort” scenario of no external mixing and no additional ART linkage in the control arm (Figure 2A), mean cumulative 3-year incidence in the control arm, averaged across all simulations, is estimated to be 1.73% compared to 1.04% in the intervention arm. The relative risk (RR) between the intervention and control arms is estimated to be 0.60 (95% confidence interval [CI] 0.54-0.67, p<0.001), with 0% of simulations yielding a nonsignificant result. In the second scenario (Figure 3B) – a closed cohort but with maximum ART linkage in the control arm – the RR is 0.86 (95% CI 0.77-0.97, p=0.014), with 38% of simulations showing a non-significant result. In the third scenario with equivalent mixing between SEARCH and non-SEARCH residents, but no additional ART linkage in the control arm, the RR is 0.83 (95% CI 0.76-0.92, p<0.001), with 18% of simulations non-significant. Finally, if equivalent external mixing and additional ART linkage in the control arm occur together (Figure 3D), the RR is 0.96 (95% CI 0.87-1.06, p=0.458), and 72% of simulations are non-significant. Across all permutations of external mixing and ART linkage in the control arm, the SEARCH Study is expected to show between a 4-40% reduction in new infections, though there is a substantial probability that the trial would show a non-significant result.

**Figure 3.**
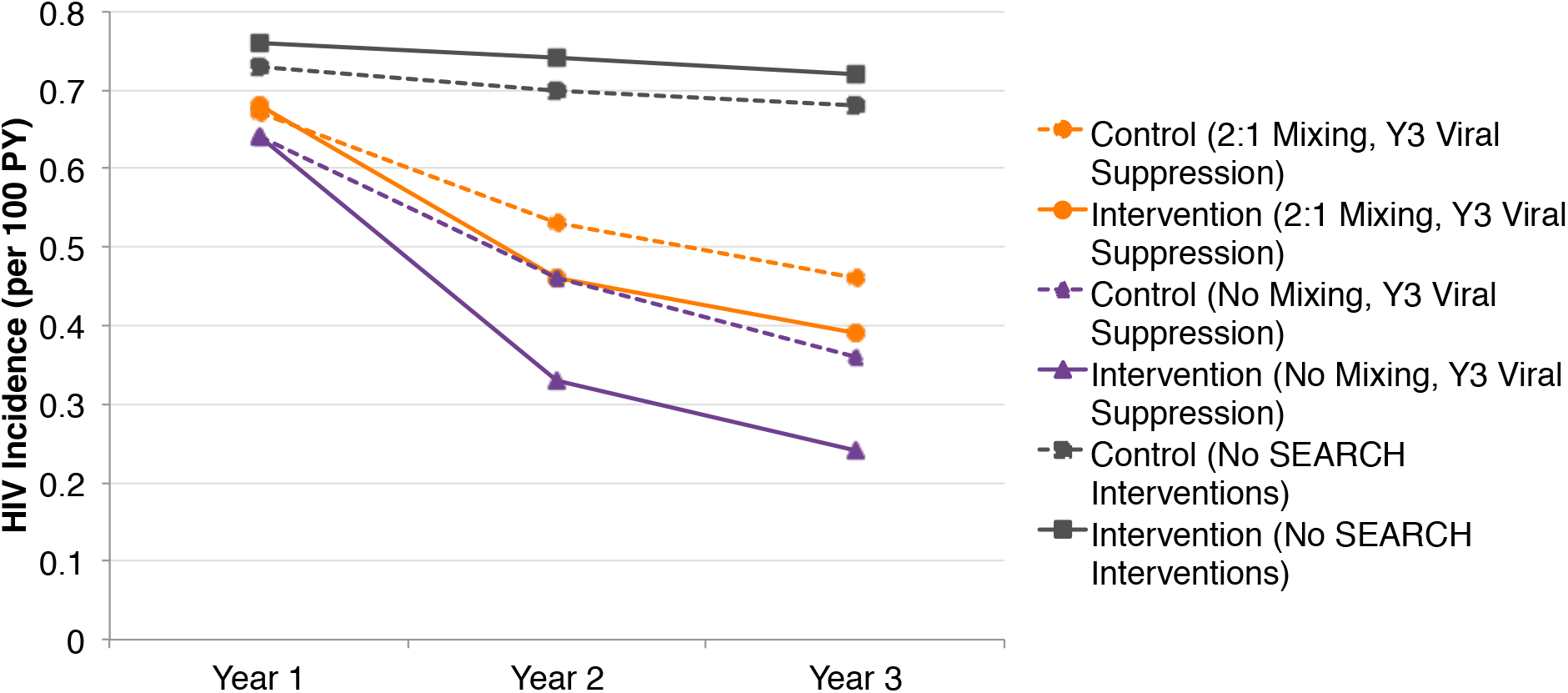
Modeled annual incidence (per 100 person-years) over time in the control (red) and intervention (blue) arms of the SEARCH trial, under three different scenarios: no SEARCH interventions in either arm of the trial (gray), no mixing and year 3 viral suppression in both arms of the trial (purple), and “best guess” assumptions using baseline mobility and year 3 viral suppression in both arms of the trial (orange). Solid lines represent the intervention arm and dashed lines represent the control arm.

In Figure 3, we additionally provide a narrower range of “best guess” scenarios based on year 3 viral suppression data in both control and intervention arms, and mobility data at baseline. The RR is estimated to be 0.90 (95% CI 0.81-1.00, p=0.05) and 49% of simulations showed a non-significant reduction in HIV incidence. In this scenario, annual incidence in the intervention arm is predicted to decline from 0.68/100 PY in the intervention arm from year 1 to 0.39/100 PY in year 3, and from 0.67/100PY in the control arm from year 1 to 0.46/100 PY in year 3. If we assume a closed cohort with no external mixing and estimate the change in incidence based on year 3 viral suppression data alone, the RR is 0.80 (95% CI 0.72-0.90, p<0.001), with 12% of simulations non-significant.

## Conclusions

The SEARCH Study represents an important opportunity to assess the population-level impact of treatment as prevention on HIV incidence. Not only has the trial exceeded the UNAIDS 90-90-90 targets, but it was also able to achieve high levels of testing, linkage, and viral suppression in populations that have historically been difficult to reach, such as men and youth [13]. In spite of these successes, multiple sources of uncertainty – namely the comparative level of ART scale-up in the control arm and the degree to which SEARCH residents mixed with non-SEARCH residents – could substantially affect the impact of the trial on HIV incidence. We estimate that the study is expected to show between a 4-40% reduction in 3-year cumulative HIV incidence due to the trial interventions, with our “best guess” prediction showing a mean 10% reduction in 3-year cumulative incidence. However, in this scenario, half of all simulations reported a non-significant difference in cumulative incidence between the two arms of the trial. Despite the known efficacy of ART in preventing HIV transmission and the achievement of the 90-90-90 targets in the SEARCH Study, the trial results are likely to not show a statistically significant reduction in new infections based on our model estimates.

Importantly, even if the trial does not show a significant result, the true population-level impact of universal test-and-treat is expected to be much larger than the impact captured in the SEARCH trial. Compared to a counterfactual with no SEARCH interventions, Modeling estimates that ART scale-up in the intervention arm – assuming no external mixing – reduced 3-year cumulative incidence from 1.78% to 1.04%, a 42% reduction. In a real-world scenario of test-and-treat, external mixing will pose less of an issue as ART coverage among surrounding communities increases.

Not only does the “active control” arm of SEARCH attenuate the measured impact of test-and-treat, but, if the difference between the arms is relatively small, it could also signal that fewer interventions are necessary to achieve linkage and viral suppression among the majority of HIV-positives. Furthermore, the cost of hybrid-mobile model of testing (a mean of $20.50 [2014 US$] per adult tested for HIV) [28] is comparable to other non-facility based methods of testing, like mobile or HBT [29], and was able to achieve close to 89% testing coverage in the first round of testing [8, 13].

There are also several model-based limitations of this analysis. First, relative risk of transmission according to viral load is not explicitly modelled; all individuals on ART in the model have a 92% reduced risk of transmission, such that no individual on ART has a 0% probability of transmitting HIV. Second, as sexual behavior data were not explicitly collected in SEARCH, such that our modelled sexual network may not represent the actual patterns of relationship formation in the SEARCH communities. In particular, relationship patterns may differ substantially between the three regions represented in the trial. Thirdly, the model does not explicitly model migration between, into, or out of nodes, though mobility may be a key element of HIV acquisition risk, although sexual mixing with partners outside the SEARCH community was implicitly modelled by stratifying the population by residency. Finally, we did not examine the potential effects of heterogeneity in suppression in this analysis. Given that approximately 20% of the HIV-positive population in the intervention arm remained unsuppressed after two years of follow-up [13], it is important to understand the characteristics and behavior of these individuals in order to be able to predict new infections accurately [21].

Our results show that, despite reaching and exceeding the 90-90-90 targets in the intervention arm, the SEARCH Study is equally likely to report a non-significant result as it is a significant result. Should the trial show no significant difference between the two arms of the trial, further work will be necessary to investigate potential explanatory factors, including mobility, heterogeneity, and an active control arm. A non-significant result may signal that HIV incidence declines can be achieved with hybrid-mobile testing alone, or that trial residents were forming a substantial portion of sexual partnerships with non-residents, while a significant result would confirm that universal test-and-treat can accomplish population-level incidence declines.

